# Tau seeding without tauopathy

**DOI:** 10.1101/2022.02.03.479049

**Authors:** Michael S. LaCroix, Brian D. Hitt, Joshua D. Beaver, Sandi-Jo Estill-Terpack, Kelly Gleason, Carol A. Tamminga, Bret M. Evers, Charles L. White, Marc I. Diamond

**Affiliations:** Center for Alzheimer’s and Neurodegenerative Diseases, Peter O’Donnell Jr. Brain Institute, University of Texas Southwestern Medical Center, Dallas, TX, USA; Department of Neurology, UT Southwestern Medical Center, Dallas, TX, USA; Department of Psychiatry, UT Southwestern Medical Center, Dallas, TX, USA; Department of Pathology, UT Southwestern Medical Center, Dallas, TX, USA

**Keywords:** Alzheimer’s disease, tauopathy, FRET biosensor, prion, tau seeding activity, healthy brain

## Abstract

Neurodegenerative tauopathies such as Alzheimer’s disease (AD) are caused by brain accumulation of tau assemblies. Evidence suggests tau functions as a prion, and cells and animals efficiently propagate unique tau assemblies. This suggests a dedicated cellular replication machinery, with normal physiologic function for tau seeds. Consequently, we hypothesized that healthy control brains would have seeding activity. We recently developed a novel monoclonal antibody (MD3.1) specific for tau seeds. We used this antibody to immunopurify tau from the parietal and cerebellar cortices of 19 healthy subjects ranging 19-65 years. We detected seeding in the parietal cortex, but not in the cerebellum, or in wild-type or human tau knockin mice, suggesting that cellular/genetic context dictates development of seed-competent tau. Seeding did not correlate with subject age or brain tau levels. Dot blot analyses revealed no AT8 immunoreactivity above background levels in parietal and cerebellar extracts and <1/100 of that present in AD. Based on binding to a panel of antibodies, the conformational characteristics of control seeds differed from AD, suggesting a unique underlying assembly, or structural ensemble. Tau’s ability to adopt self-replicating conformations under non-pathogenic conditions may reflect normal function that goes awry in disease states.

## Introduction

Accumulation of intracellular assemblies of the microtubule-associated protein tau (MAPT) underlies myriad neurodegenerative diseases termed tauopathies (Lee and Trojanowski, 1999). Alzheimer’s disease (AD), the most common tauopathy, now afflicts ∼50 million people worldwide, and is estimated to harm ∼150 million by 2050 (Guerchet et al., 2020). Most tauopathies are sporadic, while some are caused by dominantly inherited mutations in the MAPT gene (Lee and Trojanowski, 1999). The origin of sporadic tauopathies has remained elusive, but growing experimental evidence suggests that tau functions as a prion in pathological states. Initial work from our group and others suggested that tau stably propagates unique fibrillar structures *in vitro* (Frost et al., 2009b). In living systems, exogenous tau assemblies are spontaneously taken up by cultured cells, and serve as templates for intracellular aggregation (Frost et al., 2009a), and similarly tau inoculated into transgenic mouse brain induces intracellular pathology (Clavaguera et al., 2009). Diverse neuropathologies also form in a tauopathy mouse model upon brain inoculation with tau fibrils prepared from human tauopathies (Clavaguera et al., 2013). We have further observed that tau forms a variety of unique structures, or strains, that propagate indefinitely and transmit readily among cells, and, after inoculation into a mouse model, produce unique, transmissible patterns of neuropathology (Sanders et al., 2014). We also observed unique strain composition patterns in five different tauopathies, including variation within specific neuropathological diagnoses (Sanders et al., 2014). In a large survey of 18 strains propagated in cells, we determined that each gave rise to a unique pattern of neuropathology following inoculation into a mouse model (Kaufman et al., 2016). This work has been confirmed and extended by cryo-electron microscopy (cryo-EM) of tau fibril structures extracted from different tauopathies, which has revealed unique core structural features that correlate with different neuropathological diagnoses (Falcon et al., 2018; Falcon et al., 2019; Fitzpatrick et al., 2017; Zhang et al., 2020). Recent work reports fibril diversity within a single neuropathological diagnosis in multiple tauopathies (Shi et al., 2021), consistent with our prior isolation of distinct strains from individual brains (Sanders et al., 2014). Taken together, considerable evidence supports the idea that pathological assemblies of distinct structure drive the development of unique tauopathies.

The faithful maintenance of tau strains in cells, mice, and humans, suggests the existence of an intrinsic replication machinery that participates in the amplification of unique structures. The normal function of tau is not entirely clear. Tau binds microtubule filaments (Cleveland et al., 1977; Weingarten et al., 1975) through interactions across the interface of tubulin heterodimers (Kadavath et al., 2015; Kellogg et al., 2018) and has been proposed to stabilize microtubules *in vivo* (Bunker et al., 2004; Witman et al., 1976).Yet tau knockout mice are viable (Ke et al., 2012), suggesting microtubule stabilization is not an essential function, or that it is compensated for by other proteins.

An enormous literature suggests that many proteins form self-amplifying assemblies to regulate biological processes. The first examples were described in yeast (Wickner et al., 2015), and similar protein activities in mammals are now established. For example, the prion-like polymerization of MAVS and ASC proteins transduces signaling in innate immunity and inflammation based on self-propagating assemblies (Cai et al., 2014; Hou et al., 2011). In mice, the cytoplasmic polyadenylation element-binding protein (CPEB) changes from a soluble to an aggregated state that promotes translation of sequestered synaptic mRNAs, maintaining long-term potentiation (Fioriti et al., 2015; Huang et al., 2006; Pavlopoulos et al., 2011; Rayman and Kandel, 2017; Si et al., 2003). Last, in hippocampi of mice exposed to cellular stress, TIA-1, which facilitates the assembly of stress granules, forms heritable aggregates through its amyloidogenic C-terminal prion-like domain (Rayman and Kandel, 2017). Given data that tau functions as a prion in experimental systems, we hypothesized a physiological role for tau conformers that serve as templates for their own replication, i.e. “seeds.” According to this model, tau aggregation in disease might represent a normally occurring process that escapes physiologic regulation, not a *de novo* and purely pathological function. This model predicts the existence of tau seeds in healthy individuals, but until now they have eluded detection. In this study we use immunoprecipitation of brain lysate with a novel anti-tau antibody, and a highly sensitive biosensor assay (Hitt et al., 2021) to address this question.

## Results

### Tau seeding in parietal cortex

In preliminary work we screened available antibodies for those that would most efficiently precipitate seeding activity from control brain, and identified MD3.1, an antibody raised against a peptide of R1/R3 of the tau repeat domain (aa263-311 Δ275-305) with a “trans” proline residue consisting of N-Boc-trans-4-fluoro-L-proline substituted at P270. MD3.1 binds tau seeds with high efficiency (B. Hitt, *in preparation*). We then used MD3.1 to test brain lysates from 19 control subjects of diverse ages (19-65 y.o.) (**Table 1**).

**Table 1.** Demographic Data from the cases studied. The 19 tauopathy negative control cases studied span six decades of life ranging in age from 19 to 65. Abbreviations: **F/M** – female/male; **age** – age in years; **C** – Caucasian; **PI** – Pacific Islander; **CTRL** – control, **PMI** - post-mortem interval in hours.

| Case | Sex | Age | Race | Diagnoses | PMI (hrs) | Manner and Cause of Death |
| --- | --- | --- | --- | --- | --- | --- |
| A | M | 20 | C | Control | 21.2 | Accident - chest compression from blunt force injuries |
| B | M | 47 | C | Control | 24 | Natural - Cardiovascular disease and diabetes |
| C | M | 34 | C | Control | 23.4 | Natural - Cardiovascular disease and diabetes |
| D | M | 52 | C | Control | 24 | Accident - Electrocution |
| E | F | 45 | C | Control | 22.5 | Natural - cardiovascular disease |
| F | M | 29 | C | Control | 27.4 | Natural - polycystic kidney disease |
| G | F | 50 | C | Control | 23.3 | Natural - adrenocortical deficiency |
| H | F | 62 | C | Control | 19.4 | Natural - chronic alcoholism |
| I | M | 48 | C | Control | 14.7 | Natural - mitral valve regurgitation |
| J | M | 54 | C | Control | 19.3 | Natural - cardiovascular disease |
| K | M | 19 | C | Control | 20 | Homicide - gunshot wound |
| L | M | 36 | C | Control | 23 | Undetermined |
| M | M | 34 | C | Control | 24 | Accident - drowning |
| N | F | 49 | PI | Control | 15.3 | Natural - cardiovascular disease |
| O | F | 65 | C | Control | 11 | Natural - cardiovascular disease |
| P | M | 65 | C | Control | 14 | Natural - respiratory arrest |
| Q | M | 63 | C | Control | 14 | Natural - myocardial infarction |
| R | F | 55 | C | Control | 25 | Natural - intra-cerebral hemorrhage |
| S | M | 32 | C | Control | 21.5 | Natural- pulmonary embolism |

We prepared soluble protein lysates from fresh frozen tissue of the cortex of the parietal lobe, and immunoprecipitated the total soluble protein lysate using MD3.1 antibody. We then used Lipofectamine 2000 to deliver total protein lysate (T), tau-depleted IP supernatants (S), and tau-enriched IP pellets (P) into v2H tau biosensors (Hitt et al., 2021). Non-treated and Lipofectamine-treated cells were used as negative controls; recombinant heparin-derived 2N4R tau fibrils (1pM monomer equivalent) were used as a positive control. All tau-enriched IP pellets exhibited seeding activity levels beyond that of Lipofectamine-treated controls, with 16/19 tau-enriched IP pellets reaching statistical significance (p < 0.05) compared to Lipofectamine-treated controls when tested by ANOVA (**Fig. 1**). MD3.1 did not induce a transition from inert to seed competent tau when tested with recombinant tau monomer (**Fig. 1 – Supplemental**). We concluded that human cortical tissue contains tau seeding activity, in the absence of known tauopathy.

**Figure 1.**
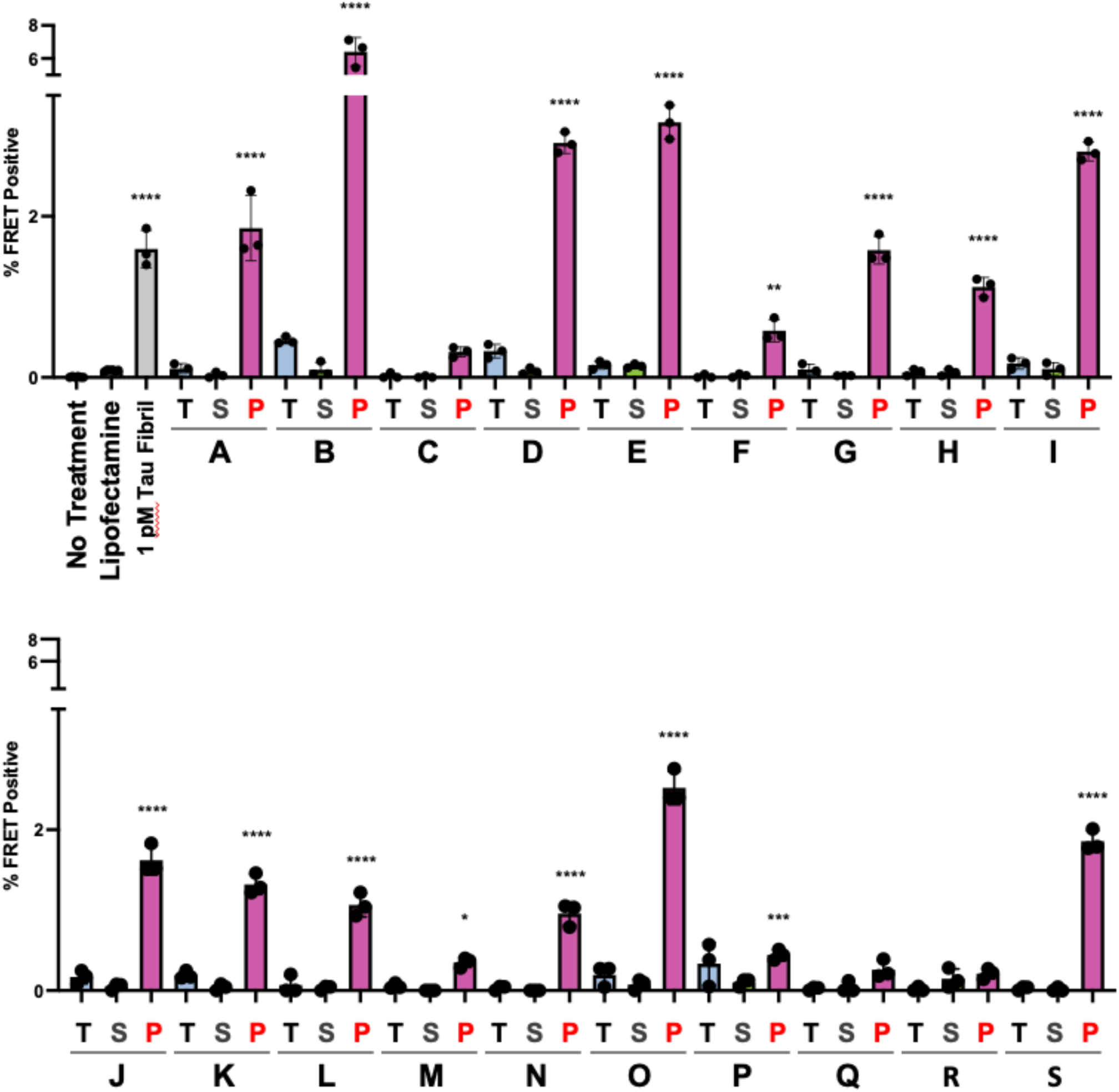
Tau seeding is present in the parietal lobe of 19 control subjects. Parietal Cortex fresh frozen samples from 19 individuals (A-S) without any known neurological diagnoses were used to create total (**T**) clarified lysate [10% (wt/vol)] followed by immunoprecipitation with the MD3.1 antibody to generate a tau depleted supernatant (**S**) and tau enriched pellet (**P**). Tau seeding was reliably detected in 16 of 19 cortical IP pellets. Columns represent the mean FRET positivity from three technical replicates (dots). Statistical significance was determined by performing one-way ANOVA followed by Dunnett’s multiple comparisons testing of all samples compared against Lipofectamine treated negative controls, *p < 0.05, **p < 0.01, ***p < 0.001, ****p < 0.0001. Errors bars = S.D.

**Figure S—Supplemental.**
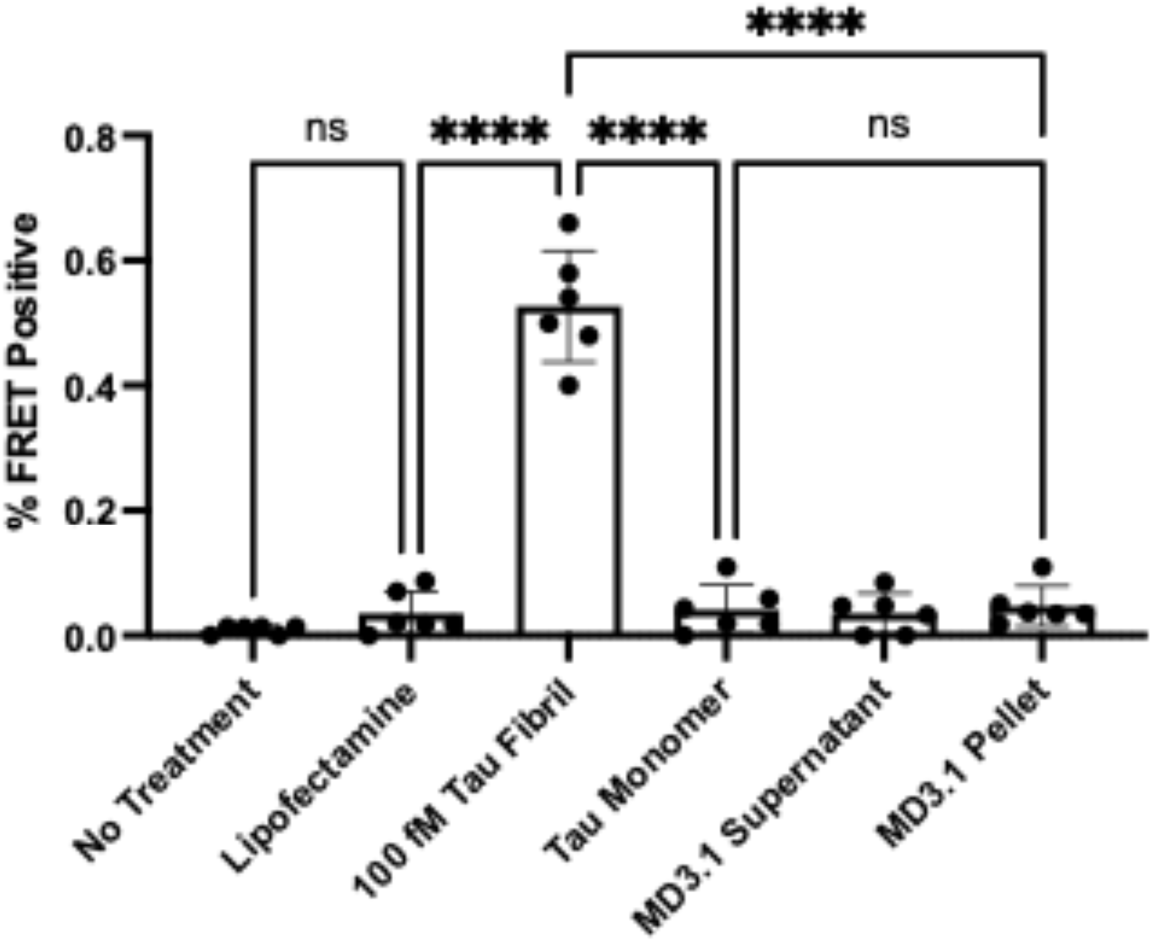
MD3.1 antibody does not induce seed conversion of recombinant tau monomer under experimental immunoprecipitation conditions. No seeding activity was detected in the pellet after immunoprecipitation of 500ng recombinant 2N4R tau with MD3.1. Statistical significance was determined by performing one-way ANOVA followed by Tukey’s multiple comparisons test, *p < 0.05, **p < 0.01, ***p < 0.001, ****p < 0.0001. Errors bars = S.D.

### Tau seeding is absent in the cerebellum

A past study found that tau seeding is largely absent from the cerebellum except at late Braak stages (Furman et al., 2017). Consequently, we tested for seeding activity in the cerebellum samples. We prepared total soluble protein lysates and performed immunoprecipitation with the MD3.1 antibody. We did not detect significant seeding in any sample (**Fig. 2**).

**Figure 2.**
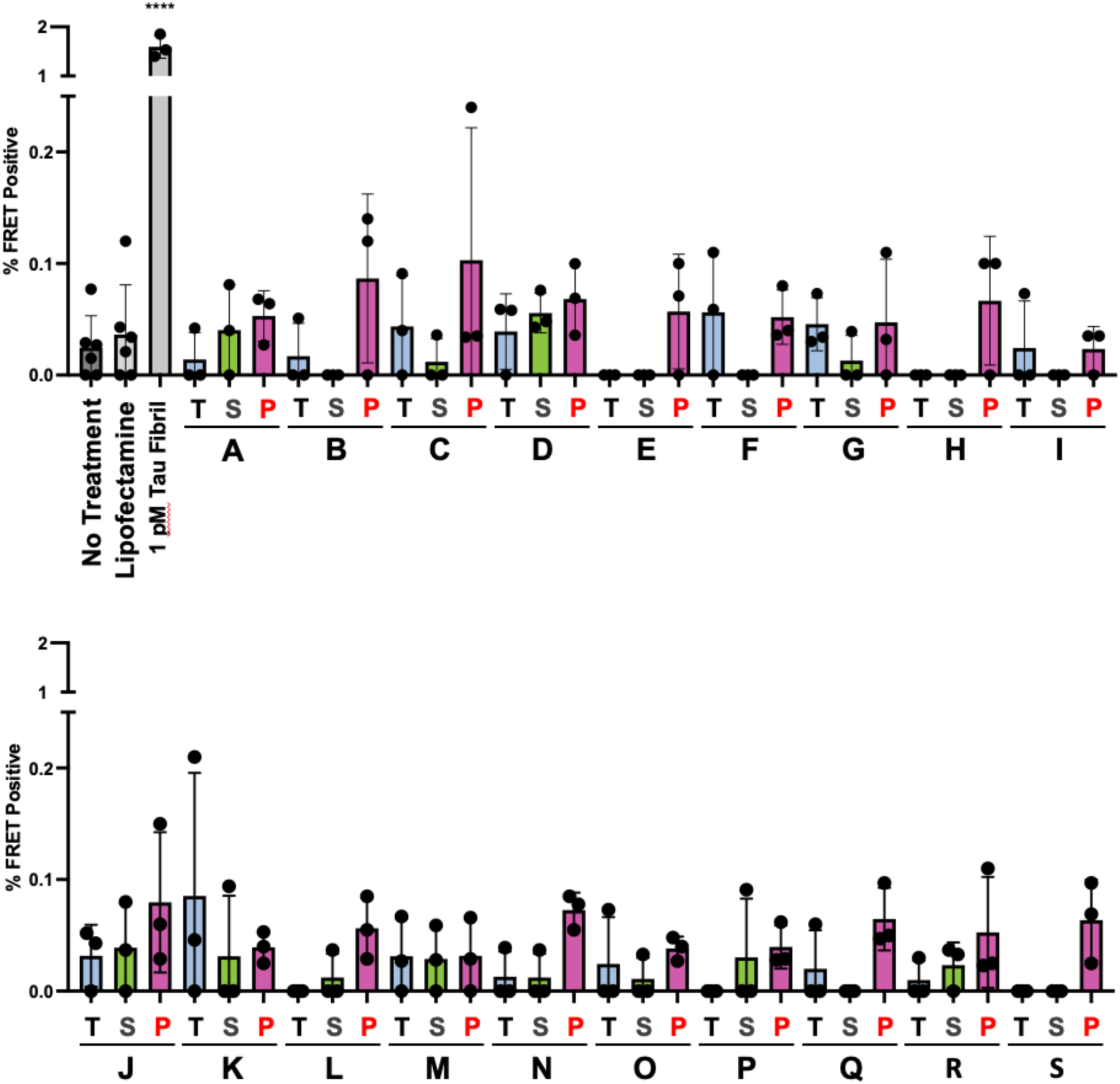
No reliable tau seeding activity is detected in the cerebellum of 19 control subjects. Fresh frozen cerebellum samples from 19 subjects without any known neurological diagnoses were used to create total (**T**) clarified lysate [10% (wt/vol)] followed by immunoprecipitation with the MD3.1 antibody to generate a tau depleted supernatant (**S**) and tau enriched pellet (**P**). No reliable tau seeding was detected across all cerebellum samples. Columns indicate the mean FRET positivity from three technical replicates (dots). Statistical significance was determined by performing one-way ANOVA followed by Dunnett’s multiple comparisons testing of all samples compared against Lipofectamine treated negative controls, *p < 0.05, **p < 0.01, ***p < 0.001, ****p < 0.0001. Errors bars = S.D.

### AT8 signal is absent in brain lysates

One explanation for our results was that we had missed underlying tauopathy in the cases we studied, despite their selection as healthy controls. Because it was impossible to carry out histopathology on the precise brain regions studied by the seeding assay, we analyzed homogenates for phospho-tau using AT8. This mouse monoclonal antibody recognizes p-Ser202 and p-Thr205, and is an accepted standard for diagnosis of tauopathy (Biernat et al., 1992). We used a dot blot analysis to be sure that any larger assemblies would be fully detected, spotting 2 µg of brain lysate on nitrocellulose membrane for detection with AT8. We used an AD brain lysate in serial dilutions for reference (**Fig. 3**). We observed signal in AD lysate at approximately 100x that observed in all parietal and cerebellar samples studied. We observed no difference in signal between parietal samples, which contained seeding activity, and cerebellar samples that did not. We concluded that there was no evidence of tauopathy within the brain tissues we analyzed.

**Figure 3.**
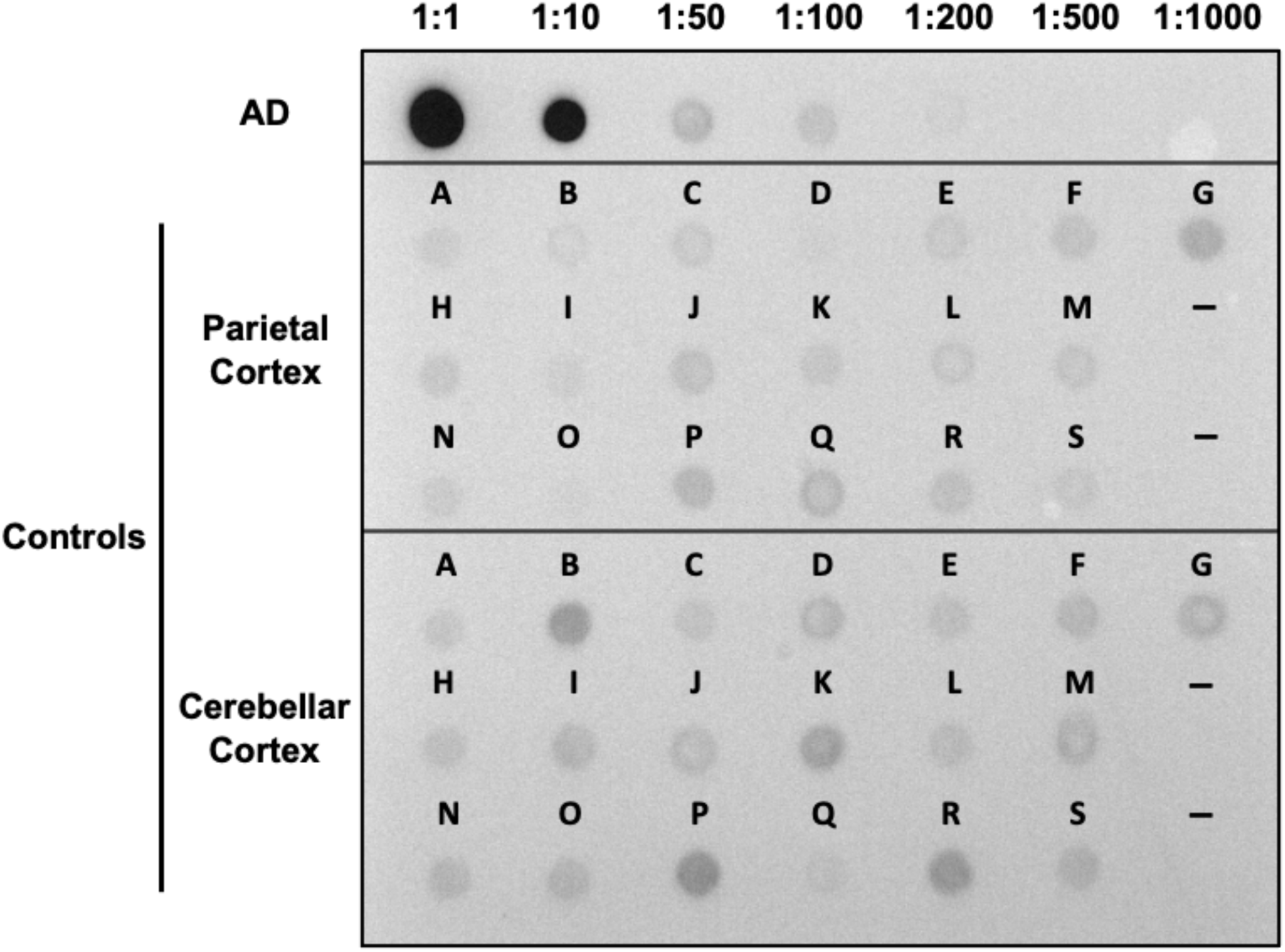
Control brains contain markedly less AT8 positivity relative to AD. 2 µg of soluble protein lysate from AD was used for 1:1, followed be serial dilutions. 2 µg total soluble protein was used for control samples, loaded in left to right order starting with sample A. Dashes represent no loading.

### Seeding is independent of total tau levels and subject age

To test the correlation of tau levels and seeding activity, we used an ELISA developed by the Davies laboratory (Acker et al., 2013). Cerebellar protein lysates contained roughly half the tau of cortical lysates, although the ranges overlapped (**Fig. 4A**). Tau concentration in the cerebellar tau-enriched pellets was ∼60% of the cortical pellets (**Fig. 4B**). These findings were consistent with previous work that indicated overall tau concentration doesn’t correlate directly with seeding activity (Furman et al., 2017).

**Figure 4.**
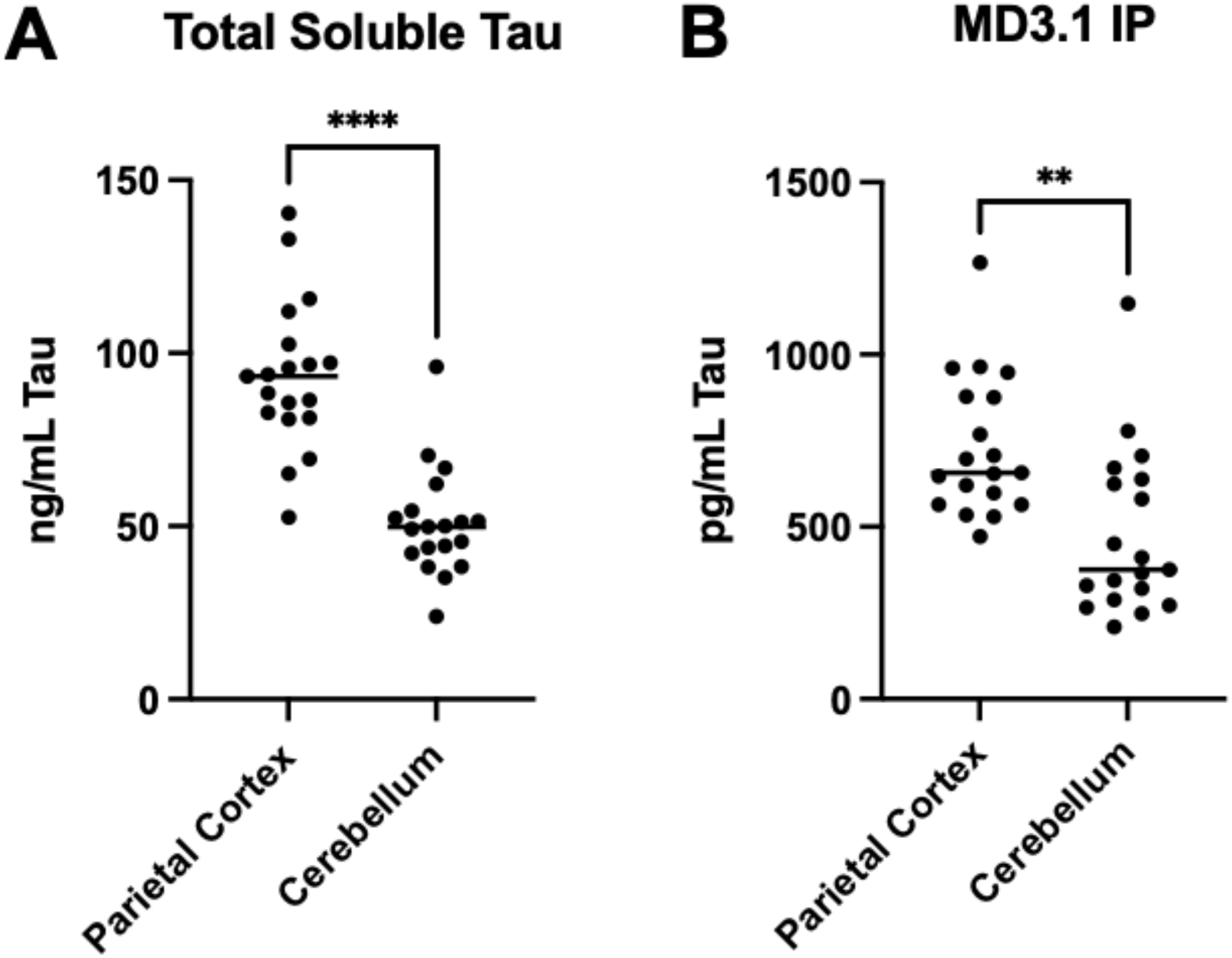
ELISA quantification of tau in brain samples. (A) Quantification of soluble tau in total clarified lysates from the parietal cortex and cerebellum. (B) Quantification of tau in the pellet following immunoprecipitation using the MD3.1 antibody. Statistical significance was determined by performing Student’s t-test, **p < 0.01, ****p < 0.0001.

We further tested the correlation of tau levels and seeding activity for the cortical samples. We observed no correlation of seeding with total tau levels (**Fig. 5A**) or with levels of tau following immunoprecipitation (**Fig. 5B**). Age is the primary risk factor for the most common tauopathy, Alzheimer’s disease, and thus we tested for its correlation with control seeding activity. We observed no correlation between seeding and age (**Fig. 5C**). We concluded that the seeding activity we observed most likely did not represent an incipient age-dependent tauopathy.

**Figure 5.**
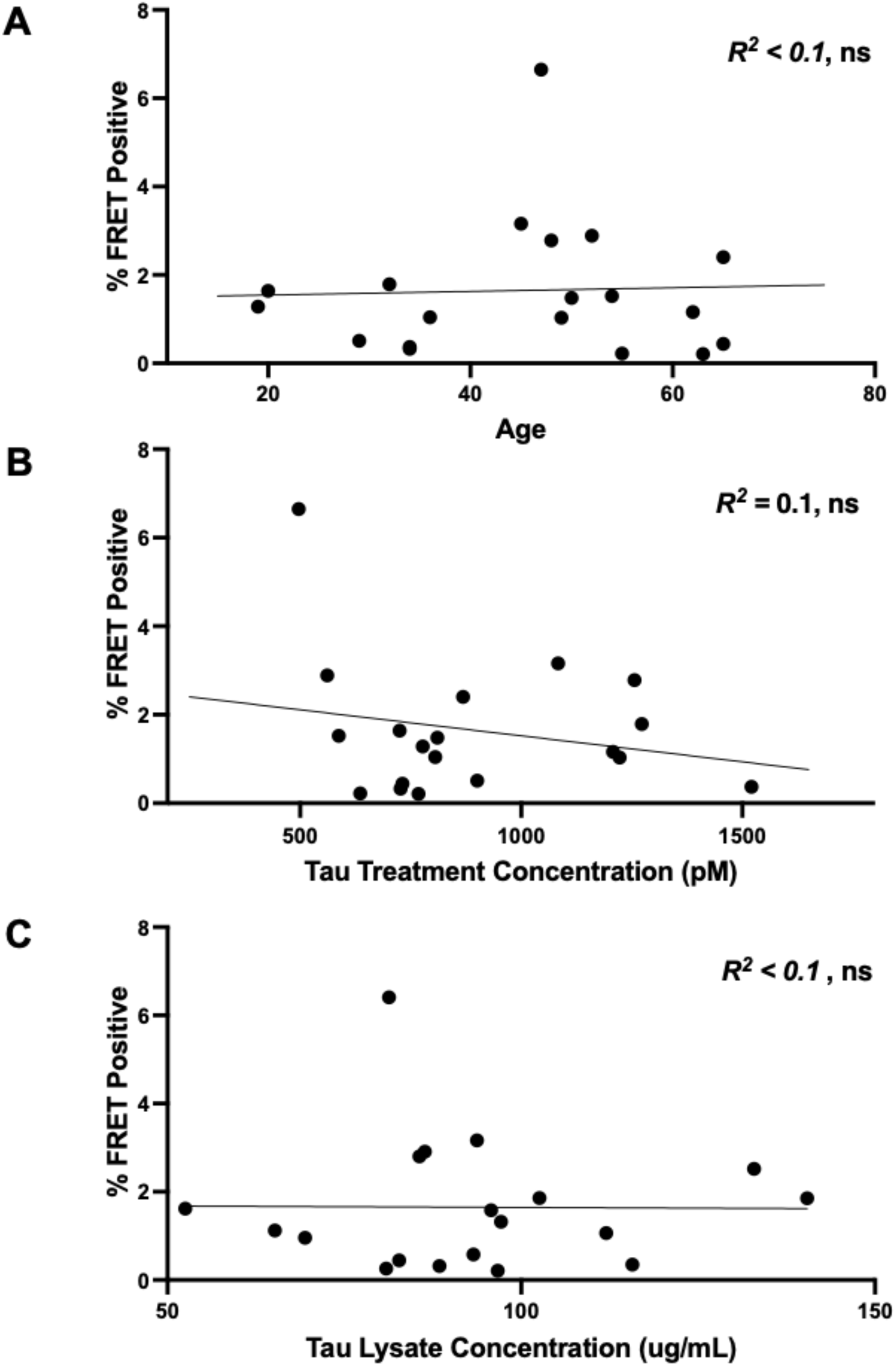
Relationships between tau seeding, age, and tau concentration in seeding samples. (A) Age did not correlate with seeding in immunoprecipitation pellets. (B) The final tau treatment concentration from immunoprecipitation pellets did not correlate with seeding in immunoprecipitation pellets. (C) Initial tau concentration in total soluble protein fractions did not correlate with seeding in immunoprecipitation pellets. Data were analyzed using Pearson correlation, ns = not significant.

### Tau seeding is absent in wild-type and hTau mice

The low or absent tau seeding in the cerebellum suggested that its development might be cell-type dependent. Human tau expressed in a knockin mouse provided the perfect opportunity to extend this inquiry. These mice (a gift from Pfizer) express all six isoforms of human tau under the mouse promoter. We first confirmed that these animals express human tau by using western blot with HJ8.5, a monoclonal antibody specific for human tau (Yanamandra et al., 2013). We prepared total protein lysates from the brains of adult human tau knock-in mice (hTau), BL6/C3H wild-type (WT), and tau knockout mice. We detected tau in knockin mouse brain with HJ8.5, which did not detect mouse tau in WT mice (**Fig. 6 – Supplemental A**). By contrast, a polyclonal anti-tau antibody (A0024, DAKO) that detects mouse and human tau revealed tau protein in hTau and WT mice (**Fig. 6 – Supplemental B**). We used immunoprecipitation with MD3.1 to test for seeding in the mouse brains, with human samples as a positive control. We detected no significant seeding activity in any mouse derived samples, including the tau-enriched IP pellet (**Fig. 6**). We concluded that tau expression in a cortical neuron is not sufficient to induce a seed-competent form, and other factors must be required.

**Figure 6.**
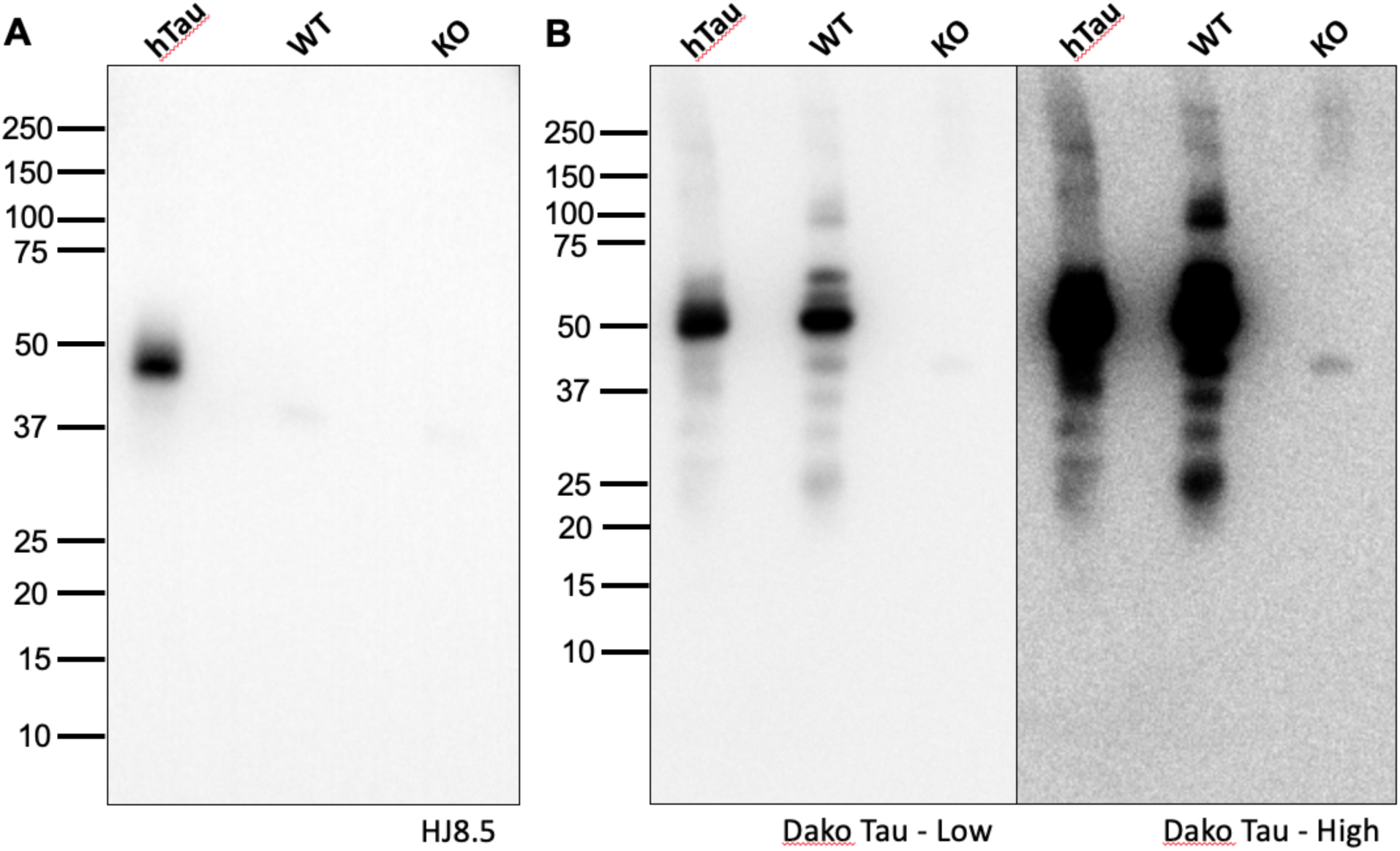
Human and mouse tau expressed in mouse cortex does not form detectable seeds. Of the groups tested, only human tau enriched via immunoprecipitation from human cortex, and not human cerebellum, formed seeds that were detectable at a statistically significant level. Tau immunoprecipitated from the cortex of mice expressing human tau (**hTau**, n=10, F=5, M=5) and wild type mouse tau (**WT**, n=9, M=4, F=5) did not show significant seeding activity. Statistical significance was determined by performing one-way ANOVA followed by Dunnett’s multiple comparisons testing of all samples compared against lipofectamine treated negative controls, ****p < 0.0001. Errors bars = S.D. Abbreviations: **Ctx –** cortex, **CB** – cerebellum.

**Figure 6– Supplemental.**
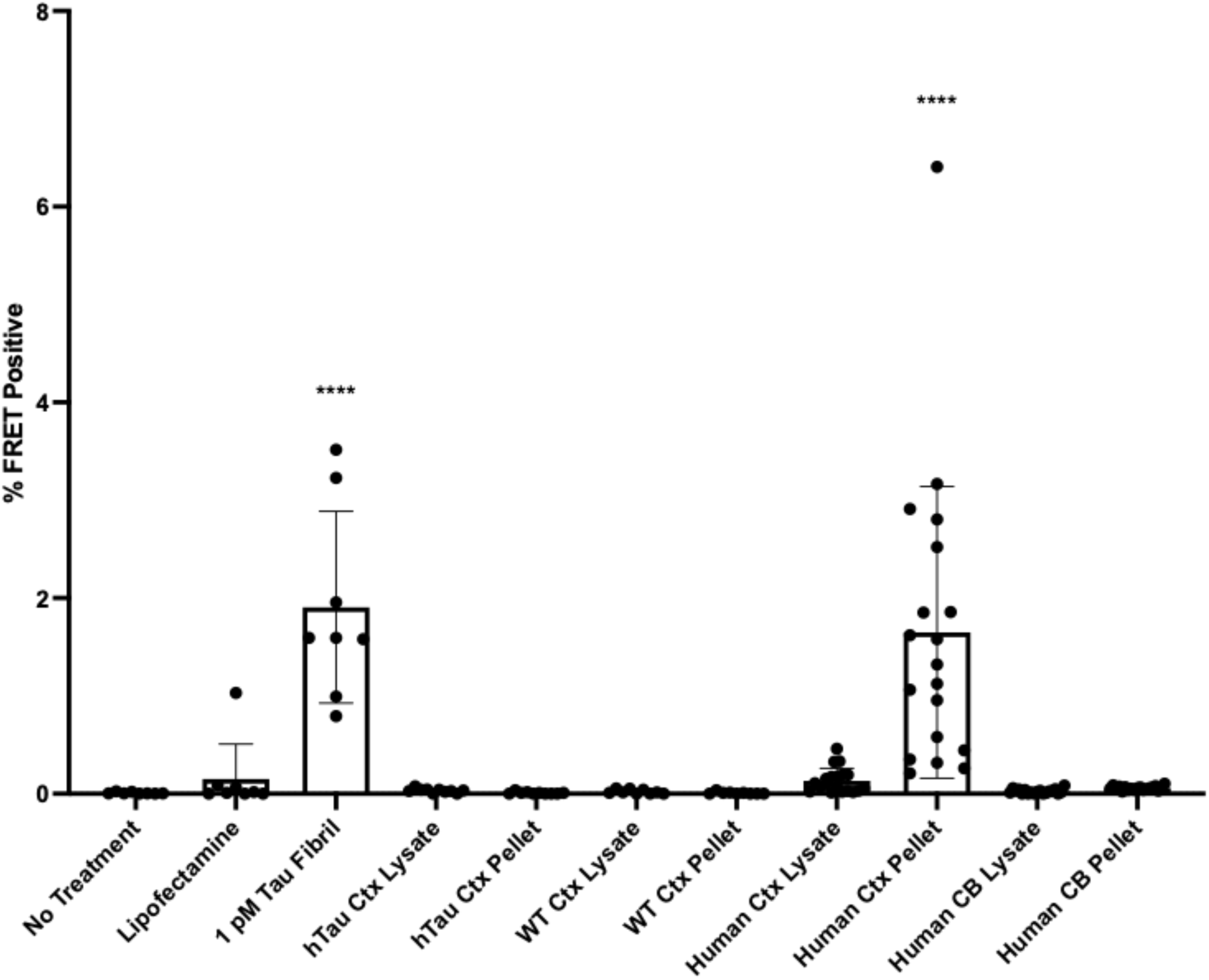
Western blots showing human tau expression in hTau mice gifted from Pfizer. (A) HJ8.5, a human tau specific antibody recognizing n-terminal residues 25-30, shows human tau expression in Pfizer human tau mice. No human tau is detected in WT mice, or tau knockout mice used as a negative control. (B) Low exposure of Dako polyclonal tau antibody reveals tau expression in hTau and WT mice, human and mouse respectively. High exposure reveals non-specific bands in knock-out mice.

### Control seeding exhibits distinct epitope accessibility vs. AD

Tau assemblies in different tauopathies exhibit conformational variation that can be resolved using cryo-electron microscopy. However, in the absence of insoluble tau it is impossible to determine the structure of the tau seeds we detected. Although the subjects we studied had no evidence of AD, we further tested the conformation of the seeds using a panel of antibodies raised against distinct epitopes across the protein (**Fig. 7A**). Based on dilution, AD brain contained ∼1,000 x more seeding activity vs. the control (**Fig. 7B**). MD3.1 most efficiently immunoprecipitated tau seeds from control brain, while antibodies against R1 and more N-terminal residues more efficiently immunoprecipitated AD seeds vs. antibodies directed against R3/R4 and more C-terminal residues (**Fig. 7C**). MD5.1, directed against residues of R3/R4, most efficiently precipitated AD seeds (**Fig. 7D**). Notably, compared to AD, MD6.2, directed against R4R’, inefficiently bound control seeds. Given the differential efficiencies in seed capture for the antibody panel, we concluded that control seeds likely represented a distinct strain, or ensemble of strains vs. AD.

**Figure 7.**
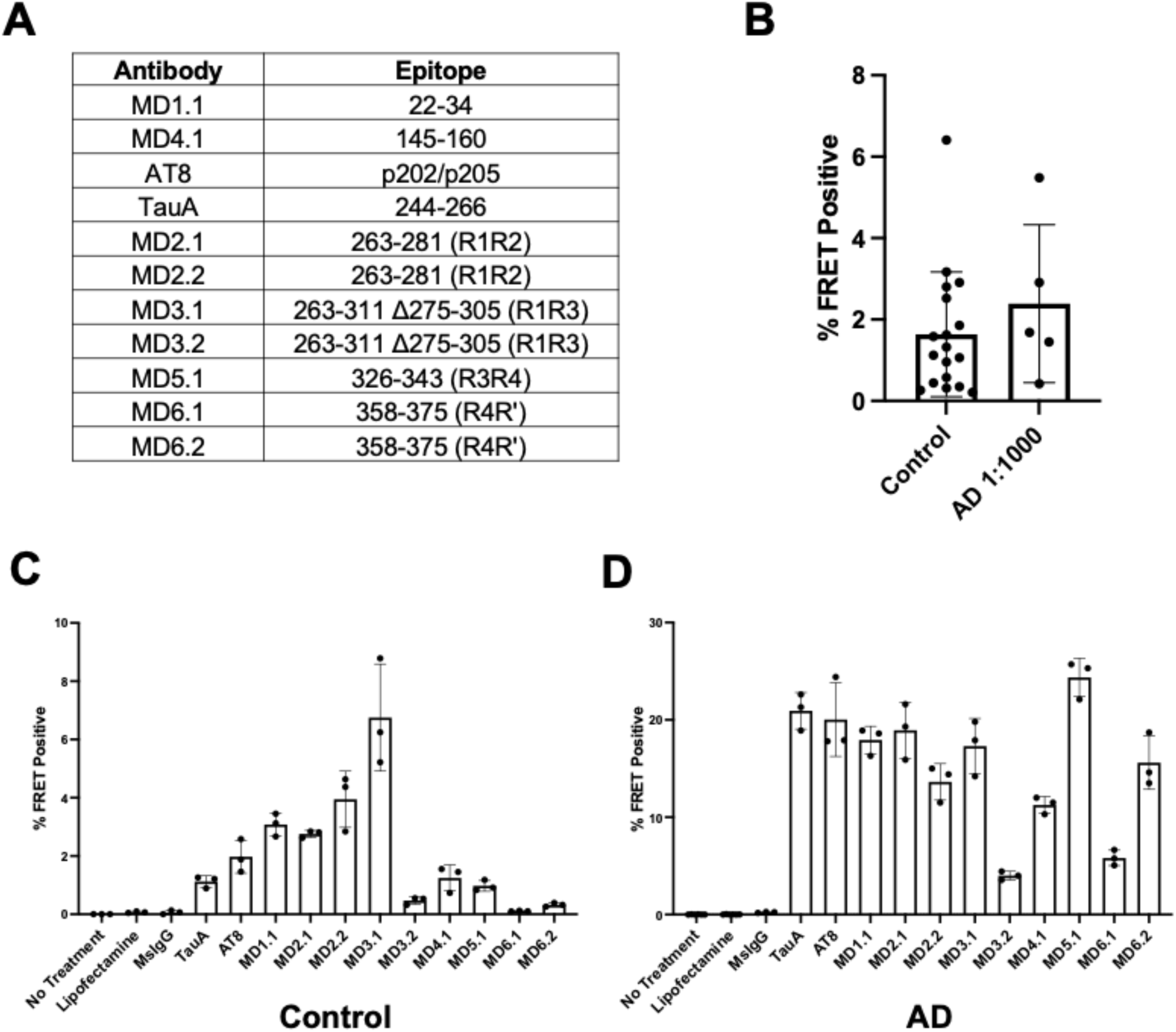
Differential seed capture efficiency from control and AD brain. A custom antibody panel reveals unique epitope exposure of tau seeds in control brain versus AD. (A) Epitopes of antibodies used. (B) MD3.1 immunoprecipitation pellets from AD brain have roughly 1000x seeding activity vs. MD3.1 pellets from control brain. MD3.1 pellets from AD were diluted 1000-fold prior to seeding on v2H biosensors while control pellets were used undiluted. No significant difference between undiluted control pellets and 1000-fold diluted AD pellets was found (p=0.3692), student’s t-test, error bars = S.D. (C) MD3.1 was most efficient at isolating tau seeds from control brain. (D) Multiple antibodies efficiently isolated seeds from AD brain.

## Discussion

In this study we have used a conformation-specific antibody and an ultra-sensitive tau biosensor cell line to detect tau seeding in the parietal cortex in 19 control individuals with no known neurodegenerative or psychiatric diseases. Notably, we detected insignificant seeding within the cerebellums of these subjects, and none in WT mice or knockin mice expressing the full human tau gene. We detected no AT8 immunoreactivity in the control brain samples. Control brain seeding activity did not correlate with tau levels or subject age. Tau seeding was present in control brain at levels <1/1000 of AD brain, and had different reactivity with a panel of anti-tau antibodies, suggesting that it was composed of tau conformers distinct from AD.

### Fidelity of tau seed detection

We have previously determined that P301S tau biosensors are specific for tau vs. other common amyloid proteins such as a-synuclein, and huntingtin (Holmes et al., 2014). Multiple controls indicate that the seeding we detected is not a result of our experimental manipulations. The MD3.1 antibody used to isolate tau seeds does not induce a transition from inert to seed competence in recombinant tau monomers. Additionally, we failed to detect strong tau seeding activity in human cerebellum or hTau knockin mice after tau immunopurification. If the experimental methods were causing seed conversion, we would expect to find seeding activity in any immunopurified sample containing FL human tau. These results suggest that tau in healthy brain adopts seed-competent conformations, and this activity is region-specific.

### Tau assemblies in normal brain function

The presence of tau seeds in healthy brain would be expected if replication of unique assembly structures is linked to a normal function of tau. The precedent for functional prions exists as a mechanism of signaling in the immune system (Cai et al., 2014; Hou et al., 2011) and at synapses (Rayman and Kandel, 2017). If amyloid formation is a critical aspect of tau’s normal biology, we would expect tau seeds to be present across all ages. Indeed, we found clear, statistically significant seeding in 16/19 samples. The absence of statistically significant seeding for 3 cases might relate to particular sub-regions of cortex that were sampled. In contrast to individuals with tauopathy, age did not predict seeding. For example, the youngest individual with significant seeding was 19 y.o., far too young to exhibit a sporadic tauopathy. Given the general paucity of healthy pediatric brain tissue we could not determine how early in human life tau seeds appear. It remains to be determined what is the functional role of tau seeds in healthy brain, although recent work from our lab suggests it might relate to RNA regulation (Zwierzchowski-Zarate, *Biorxiv 2022*). Indeed, multiple prior studies have indicated that RNA can induce tau seeding (Kampers et al., 1996), and that seeds are associated with neurofibrillary tangles (Ginsberg et al., 1997).

### Regional and species specificity of tau assembly formation

We observed a regional and species specificity for tau seeding. We detected seeding in the parietal cortex while observing minimal amounts in the cerebellum, and none in hTau knockin mouse brain. The absence of significant seeding in the cerebellum or an hTau mouse could reflect that tau forms no functional assemblies in these conditions, or conversely that the strains formed were not efficiently precipitated by MD3.1. This will require further study, and mechanisms controlling the formation and dissolution of tau seeds in the healthy brain will be critical to understand. If this is dynamic, then it seems likely that human genetic factors, and even the state of brain function might regulate seed abundance. Tau ligands or post-translational modifications might also regulate seed abundance. Pathogenic tau strains may thus represent perturbation of tau’s normal function, in which strains form that are inefficiently cleared, or that set in motion metabolic or genetic changes that lead to feed-forward loops of amplification.

### Control brain harbors unique tau strains compared to AD

One explanation for our findings is that control brain seeding simply represents the earliest form of AD or some other tauopathy. Because of the nature of the brain preparation in these samples, we could not directly evaluate by immunohistochemistry the material used in the seeding assays. We feel that undiagnosed tauopathy is highly unlikely here for several reasons. First, the cell-based biosensor assay we used is enormously sensitive to incipient tauopathy, and indeed predicts pathology in humans or mice prior to the development of immunohistochemical evidence of disease using AT8 or similar antibodies (Furman et al., 2017; Holmes et al., 2014). Given that none of the samples studied showed any seeding without immuno-enrichment, we feel very confident that there was no undiagnosed tauopathy in the brain samples. Second, we observed no age-dependence of control brain seeding, which would have been expected if it simply represented incipient sporadic tauopathy. Indeed, we observed relatively strong seeding in a 19 y.o. subject, an age at which it would be essentially inconceivable to observe sporadic tauopathy. Third, we directly evaluated brain lysates using dot blot, to capture both aggregated and non-aggregated forms of tau. We detected similar background AT8 signal in conditions where we observed seeding (parietal cortex) and where we did not (cerebellar cortex), in both cases at a level <1/100 of that detected in AD. Finally, using antibodies that target epitopes across tau, we observed different immunoprecipitation efficiencies for control vs. AD seeds. It is possible that post-translation modifications or associated cofactors led to differential immunoprecipitation, with the underlying seed conformation being unchanged. However, the simplest interpretation of our findings is that control seeds differ in epitope exposure because of their assembly structure, i.e. strain identity.

In summary, we have detected tau seeds by immunoprecipitation in healthy brain in an age-independent, region-specific manner. The presence of seeds in healthy individuals suggests that tau forms self-replicating assemblies that may play a physiological role, rather than being purely pathologic. We predict that cofactors regulate the transition from inert to seed competent tau, and we hope to identify them in future work. In particular, more research must elucidate whether tau seeds have a functional role in healthy brain, and if critical co-factors or post-translational modifications lead to dysregulated tau assembly formation. This could be especially important in light of therapies to clear tau assemblies, or inhibit their formation, which might have unintended consequences for normal brain function.

## Materials and Methods

### Biosensor cell line v2H

Highly sensitive second-generation tau biosensor cells termed v2H (Hitt et al., 2021) were used for seeding assays. These cells are based on expression of tau repeat domain fragment (246-378) containing the disease-associated P301S mutation (tau-RD) fused to mCerulean3 or mClover3. The v2H line was selected for high expression with low background signal and high sensitivity. Seeding experiments used previously established protocols (Holmes et al., 2014).

### Cell culture

v2H biosensors were grown in Dulbecco’s Modified Eagle’s medium (Gibco) supplemented with 10% fetal bovine serum (HyClone), and 1% glutamax (Gibco). For terminal experiments, 1% penicillin/streptomycin (Gibco) was included. Cells were tested free of mycoplasma (VenorGem, Sigma) and cultured at 37°C with 5% CO_2_ in a humidified incubator. To avoid false-positive signal from v2H biosensors, cells were passaged prior to ∼80% confluency.

### Human brain samples

Human brain tissue was obtained from 19 control subjects (6 females, 13 males, age range 19-65 years, Table 1) without any known tauopathy or psychiatric diagnoses, with Institutional Review Board (IRB) approval at University of Texas Southwestern Medical Center. Informed written consent for donation of tissue was obtained from next of kin prior to collection. Brains were sectioned and flash frozen in liquid nitrogen for long-term storage at -80°C. Pulverized frozen tissue from the cortex of the parietal lobe and cerebellum was used to prepare total soluble protein lysates for further experiments.

### Human sample preparation

Fresh frozen pulverized tissue was suspended in tris-buffered saline (TBS) containing cOmplete mini protease inhibitor tablet (Roche) at a concentration of 10% w/vol. Samples were then dounce homogenized, followed by pulsing probe sonication at 75 watts for 10 min (Q700, QSonica) on ice in a hood. The sonication probe was washed with a sequence of ethanol, bleach, and distilled water to prevent cross-contamination. Lysates were then centrifuged at 23,000 x g for 30 min and the supernatant was retained as the total soluble protein lysate. Protein concentration was measured with the BCA assay (Pierce). Fractions were aliquoted and stored at -80°C prior to immunoprecipitation and seeding experiments.

### Mouse lines

Tau KO mice (n=10, 5 females, 5 males, average age 12.3 months) containing a GFP-encoding cDNA integrated into exon 1 of the MAPT gene were negative controls (Tucker et al., 2001). Tau KO mice were obtained from Jackson Laboratory and maintained on a C57BL/6J background. Wild-type mice (n=9, 5 females, 4 males, average age 12.9 months) of BL6/C3H background were used as a source of murine tau. We used mice on BL6 background with the human MAPT gene knocked in at the mouse tau locus that express all six isoforms of human tau (n=10, 5 females, 5 males, average age 16.2 months) as a source of murine-expressed human tau (gift from Pfizer), and verified by genomic sequencing in the Diamond lab. All mice involved in this study were housed under a 12 hour light/dark cycle, and were provided food and water *ad libitum*. All experiments involving animals were approved by the University of Texas Southwestern Medical Center Institutional Animal Care and Use Committee (IACUC).

### Mouse sample collection and preparation

Mice were anesthetized with isoflurane and perfused with chilled phosphate buffered saline (PBS) + 0.03% heparin. The forebrain and cerebellum were separated and weighed prior to flash freezing in liquid nitrogen and storage at -80°C. As described previously for human tissues, fresh frozen forebrain was suspended in TBS containing cOmplete mini protease inhibitor tablet (Roche) at a concentration of 10% w/vol. Samples were then dounce homogenized, followed by pulsing probe sonication at 75 watts for 10 min on ice in a hood (Q700, QSonica). The sonication probe was washed with a sequence of ethanol, bleach, and distilled water to prevent cross-contamination of seeding activity. Lysates were then centrifuged at 23,000 x g for 30 min and the supernatant was retained as the total soluble protein lysate. Protein concentration was measured with the BCA assay (Pierce). Fractions were aliquoted and stored at -80°C prior to immunoprecipitation and seeding experiments.

### Immunoprecipitation from protein lysate or tau monomer

Immunoprecipitations were performed using 50 µL of magnetic Protein A Dynabead slurry (Thermofisher), washed twice with immunoprecipitation (IP) wash buffer (0.05% Triton-X100 in PBS), followed by a 1 hour room temperature incubation with 20 µg of anti-tau antibody. Beads were washed three times in IP wash buffer, 1000 µg of total protein lysate or 500 ng of recombinant tau monomer was added to the Protein A/anti-tau antibody complexes on the beads and rotated overnight at 4 °C. After overnight incubation, supernatant was removed as the tau-depleted fraction and the beads were washed three times in IP wash buffer and then moved to clean tubes for elution. IP wash buffer was removed and beads were then incubated in 65 µL of IgG Elution Buffer (Pierce) for 7 min to elute tau. The elution buffer was collected in a separate microcentrifuge tube and a second elution step in 35 µL of IP elution buffer was performed for 5 min and pooled with the initial elution. The elution was then neutralized with 10 µL of Tris-HCl pH 8.4 to finalize the tau-enriched IP pellet.

### Enzyme-linked immunosorbent assay for total tau

A total tau “sandwich” ELISA was performed as described previously (Acker et al., 2013). Anti-tau antibody reagents were kindly provided by the late Dr. Peter Davies (Albert Einstein College of Medicine). 96-well round-bottom plates (Corning) were coated for 48 hours at 4°C with DA-31 (aa 150-190) diluted in sodium bicarbonate buffer (6 µg/mL). Plates were rinsed with PBS three times, blocked for 2 hours at room temperature with Starting Block (Pierce), and rinsed with PBS five additional times. Total protein was diluted 1:1000 in SuperBlock solution (Pierce; 20% SuperBlock in TBS) and 50 µL sample was added per well. Tau-enriched IP pellets were diluted 1:100 in SuperBlock solution and 50 µL sample was added per well. DA-9 (raised against aa 102-150) was conjugated to HRP using the Lightning-Link HRP Conjugation Kit (Innova Biosciences), diluted 1:50 in superblock solution, and 50 µL was added per well (15 µg/mL). Sample and detection antibody complexes were incubated overnight at 4°C. Plates were then washed with PBS nine times with a 15 second incubation between each wash, and 75 µL 1-Step Ultra TMB Substrate Solution (Pierce) was added. Plates were developed for 30 min and the reaction was quenched with 2 M sulfuric acid. Absorbance was measured at 450 nm using an Epoch plate reader (BioTek). Each plate contained a standard curve, and all samples were run in duplicate. Tau concentration was calculated using MyAssays four parameter logistic curve fitting tool.

### Dot blot assays

Soluble protein lysates were prepared as described previously. Lysates were diluted to a final total soluble protein concentration of 1 mg/ml in TBS containing cOmplete mini protease inhibitor tablet (Roche). For AD, serial dilutions were subsequently prepared. 2 µl/sample was spotted onto nitrocellulose membrane (ThermoFisher Scientific) and then blocked in StartingBlock (TBS) blocking buffer (ThermoFisher Scientific) for 30 min at room temperature. After blocking, the blot was incubated overnight at 4°C with AT8 primary antibody (Invitrogen) diluted 1:1000 in blocking buffer. The membrane was washed in TBS-T five times for 5 min followed by 1 hour RT incubation with peroxidase-conjugated anti-mouse secondary antibody (Jackson ImmunoResearch) diluted 1:4000 in blocking buffer. The membrane was then washed an additional five times in TBS-T. The membrane was revealed with ECL Prime western blot detection kit (Cytiva) and imaged with a Syngene digital imager.

### Transduction of biosensor cell lines, flow cytometry and seeding analyses

The seeding assay was conducted as previously described (Holmes et al., 2014) with the following changes: v2H cells were plated 20 hours before seed transduction at a density of 16,000 cells/well in a 96-well plate in a media volume of 180 µL per well. Mouse and human total protein lysates were thawed on ice, while tau-depleted IP supernatants and tau-enriched IP pellets were isolated just before seeding. For total protein lysates and tau-depleted supernatants 10 µg of protein was used per well. For tau-enriched pellets, 10 µL of elution was used per well. Samples were incubated for 30 min with 0.5 µL Lipofectamine 2000 (Invitrogen) and OptiMEM such that the total treatment volume was 20 µL. For each experiment, cells treated with OptiMEM alone and Lipofectamine 2000 in OptiMEM were used as negative controls. The v2H line, which expresses high levels of tau RD, can show false-positive FRET signal when treated with Lipofectamine 2000, which is mitigated by passaging prior to ∼80% confluency. Recombinant tau fibrils at 1 pM and 100 fM (monomer equivalent) were used for positive controls. Cells were incubated for an additional 48 hours after treatment prior to harvesting. Cells were harvested with 0.25% trypsin and fixed in 4% PFA for 10 min, then resuspended in flow cytometry buffer (HBSS plus 1% FBS and 1 mM EDTA). The LSRFortessa SORB (BD Biosciences) was used to perform FRET flow cytometry. Single cells double-positive for mCerulean and mClover were identified and the % FRET positive cells within this population was quantified following a gating strategy previously described (Holmes et al., 2014). For each experiment 10,000 cells were analyzed in triplicate. Flow data analysis was performed using FlowJo v10 software (Treestar).

### Statistical analyses

Coded samples were obtained by M.S.L. from the lab of C.A.T. M.S.L. remained blinded prior to all seeding analyses. Flow cytometry gating and analysis of seeding activity was completed prior to decoding and interpreting the results. All statistical analysis was performed using GraphPad Prism v9.2.0 for Mac OS and Excel v16.52 (Microsoft).

